# *Cannabis sativa* (Hemp) seed-derived peptides WVYY and PSLPA modulate the Nrf2 signaling pathway in human keratinocytes

**DOI:** 10.1101/2024.01.26.577509

**Authors:** Euihyun Kim, Jihyeon Jang, Hyo Hyun Seo, Jeong Hun Lee, Sang Hyun Moh

## Abstract

*Cannabis sativa* (Hemp) seeds are used widely for cosmetic and therapeutic applications, and contain peptides with substantial therapeutic potential. Two key peptides, WVYY and PSLPA, extracted from hemp seed proteins were the focal points of this study. These peptides have emerged as pivotal contributors to the various biological effects of hemp seed extracts. Consistently, in the present study, the biological effects of WVYY and PSLPA were explored. We confirmed that both WVYY and PSLPA exert antioxidant and antibacterial effects and promote wound healing. We hypothesized the involvement of the nuclear factor erythroid 2– related factor 2 (Nrf2) signaling pathway in these observed effects, given that Nrf2 is reported to be a central player in the regulation of these observed effects. Molecular-level investigations unequivocally confirmed the role of the Nrf2 signaling pathway in the observed effects of WVYY and PSLPA, specifically their antioxidant effects. Our study highlights the therapeutic potential of hemp seed-derived peptides WVYY and PSLPA, particularly with respect to their antioxidant effects, and provides a nuanced understanding of their effects. Further, our findings can facilitate the investigation of targeted therapeutic applications and also underscore the broader significance of hemp extracts in biological contexts.

## Introduction

Substances derived from plants are utilized by humans for both cosmetic and therapeutic purposes across the globe, and oftentimes these substances trace their roots to traditional folk remedies. Several ongoing studies are investigating plant-derived substances, including plant extracts, phytochemical compounds, plant-derived peptides, among others, for various applications both *in vitro* and *in vivo*. Specifically, plant-derived peptides are primarily involved in plant growth and defense mechanisms through regulation of cell–cell signaling [1]. In particular, the effects of plant-derived peptides have been investigated in immortalized human cells [2], keratinocytes [3–5], dermal fibroblasts [6, 7], and periodontal ligament stem cells [8].

Hemp (*Cannabis sativa*) seeds have been investigated for various applications. In particular, hemp seeds are widely used in traditional remedies, various household and cleansing products, as well as in the production of edibles. Specifically, hemp seed oil is known for its positive effects on human nutrition and cosmetics because of the presence of linoleic and linolenic acids [9, 10].

Hemp seeds also exhibit antioxidant properties. Cattaneo et al. reported that minerals, such as potassium, magnesium, and calcium are found in hemp seeds, alongside phytochemicals with antioxidant activity, such as tocopherols and lignan derivatives [11] Additionally, Giacomo et al. reported the antioxidant and anti-inflammatory effects of hemp water extract on mouse fibroblasts and keratinocytes via metabolomic profiling [12]. Similarly, there are numerous reports on the antioxidant or other beneficial effects of hemp derivatives, such as extracts and oil, especially on the skin, and for cosmetic applications [13–16].

Girgih et al. characterized the structure and antioxidant and antihypertensive effects of short peptides derived from hemp seed protein through enzymatic hydrolysis. The group observed the highest antioxidant effects with the short peptides PSLPA and WVYY [17]. Such studies emphasize the extraction of peptides from hemp-derived sources as well as the characterization of the bioactive components of the extract that play a pivotal role in antioxidant effects [17, 18], However, the effects of hemp-derived peptides on antioxidant mechanisms have not been explored extensively. Some studies have suggested that peptides derived from various dietary or biological sources, specifically short peptides, possess various biological effects, including antioxidant [19], wound healing [19], anti-inflammatory [20], and antibacterial [21, 22], effects. The antioxidant effects of these small peptides have been suggested to be attributed to their structure; however, the precise correlation between the observed antioxidant effects and the structure of the peptides remains poorly understood. Further, the specific amino acids in the peptide are also thought to play a significant role in their biological effects, especially antioxidant effects [23–26]. Hence, the antioxidant effects of short peptides can be attributed to various factors. We speculated that the nuclear factor erythroid 2-related factor 2 (Nrf2) signaling pathway may play a role in the same as several studies have suggested that peptides derived from various sources, including hemp, are regulated by the Nrf2 signaling pathway [27–29]. In its basal state, Nrf2 is ubiquitinated by Kelch-like ECH-associated protein 1 (Keap1), preventing the translocation of Nrf2 into the nucleus [30]. Conversely, under oxidative stress, Nrf2 is released from the Keap1–Cullin 3 (Cul3)–Rbx complex, which facilitates its nuclear translocation [31]. In the nucleus, Nrf2 initiates the transcription of genes containing the antioxidant responsive element, including genes encoding catalase (CAT), heme oxygenase-1 (HO-1), superoxide dismutase (SOD), and NAD(P)H quinone oxidoreductase 1 [32, 33]. In addition to its role in the antioxidant response, the Nrf2 signaling pathway is also involved in inflammation, wound healing, immune response to bacterial infections [34–36].

Based on these findings, we hypothesized that hemp seed-derived peptides (HSDPs) directly regulate the Nrf2 signaling pathway, thereby enhancing its effects. Consistently, the present study aimed to ascertain the effects of HSDPs and to delineate their role in the well-documented antioxidant properties of hemp seed extracts.

## 2. Materials and Methods

### 2.1. Chemicals and reagents

All chemicals and reagents used in this study were obtained from Sigma-Aldrich Chemical Company unless indicated. In the case of HSDP, PSLPA and WVYY sequences were designed and proceeded with custom manufacturing through WELLPEP Co., Ltd. After diluting it in ethyl alcohol to a concentration of 10mM, Cell treatment was conducted to maintain the final alcohol concentration below 1%.

### 2.2. Culture of human keratinocyte cells and HSDP treatment

Human HaCaT keratinocytes, sourced from the American Type Culture Collection (ATCC, Manassas, VA, USA), underwent culture in dermal cell basal medium. This medium was enriched with 10% heat-inactivated fetal bovine serum (FBS) from Gibco (Carlsbad, CA, USA) and a penicillin/streptomycin mixture (100 U/mL) from Lonza (Walkersville, MD, USA). Cultivation occurred at 37°C in a humidified atmosphere infused with 5% CO2. Upon achieving 80–90% confluence, the cells were maintained in the same medium. Subsequently, the HaCaT cells were exposed to HSDP, an extract derived from roselle callus, at concentrations of 1%, 5%, and 10% for 24 hours. The experimental setup involved a control group treated with distilled water.

### 2.3. CCK-8 Assay

The study aimed to evaluate how HSDP affected the growth, propagation, and survival of human skin cells (HaCaT) using a CCK-8 assay. HaCaT cells were initially seeded at a density of 5 × 10^4^ cells per well in a 96-well plate and left to incubate for 24 hours. Following this, the cells were exposed to varying final concentrations (1%, 5%, and 10%) of HSCE for 24 hours, with sterile water utilized as the control. Post HSCE treatment, 1X CCK-8 solution (Cat. No. CCK-3000, Donginbio, Seoul, Korea) was introduced to each well, and the cells were further incubated for 3 hours. Measurement of wavelength absorbance at 450 nm was carried out using a Thermo Scientific Multiskan GO Microplate Spectrophotometer (Fisher Scientific Ltd., Vantaa, Finland). The percentage of cell viability was computed using the formula: (absorbance of treated cells / absorbance of control cells) x 100.

### 2.4. Melanin content assay

B16F1 cells (a mouse melanoma cell line) were initially seeded at a density of 1×10^5^ cells per well in 6-well plates and left to incubate under cell culture conditions for 24 hours. Following this, the cells received treatment with HSDP and continued to culture for an additional 72 hours. To stimulate melanin production, a positive control of 100 ppm Kojic acid was added, alongside 1 μM α-MSH to each sample. Post-incubation, the upper layer of the medium was removed, and cell harvesting was performed using 1X Trypsin-EDTA. The cell pellet was then treated with a 300 µL solution of 1 N NaOH, enabling the dissolution of melanin through boiling at 100°C for 30 minutes. Subsequently, a portion of the dissolved melanin solution was transferred to a 96-well plate for absorbance measurement at 405 nm. The absorbance value of each sample was quantified based on the protein amount, determining the rate of inhibition of melanin production.

### 2.5. Wound healing Assay

Culture inserts for wound healing assays (CBA-120, Cell Biolabs, USA) were added to each well of a 24-well plate. HaCaT cells were seeded inside the insert at a density of 1x10^6^ cells/well, ensuring 90% confluence, and cultured for 24 hours. Subsequently, the inserts were removed, capturing microscopic images of the cell morphology at 0 hours, followed by treatment with 100 ng/mL Epidermal Growth Factor for both samples and positive control groups. After an additional 18-hour incubation, the media was removed, and 4% PFA was added, allowing it to react for 15 minutes at room temperature for fixation. Post-fixation, the cells were washed three times with PBS and microscopic images of the cell morphology were captured at 18 hours. The extent of wound healing was quantified by measuring the cell migration area between the images taken at the time of post-insert removal after additional incubation and those taken at 0 hours using the ImageJ software.

### 2.6. Assessment of UV Protection

Keratinocytes are seeded at a density of 5x10^4^ cells/well in a 96-well plate and cultured. After 24 hours, the medium is replaced with serum-free medium for a 4-hour starvation period. Following this, the medium is removed, and the cells are exposed to UVB using the UVP® CL-1000® Ultraviolet Crosslinker equipment at a dose of 5-15 mJ/cm^2^. Subsequently, the cells are treated with HSDP and further incubated for 24 hours. A CCK-8 assay is then performed to assess changes in cell viability.

### 2.7. 1,1-Diphenyl-2-picryhydrazyl (DPPH) assay

A 0.1 mM DPPH solution of 0.5 mL is mixed with 0.4 mL of ethanol, and the required volume of HCDP is added to achieve the necessary concentration, then supplemented with distilled water to reach a total volume of 1 mL. Positive control groups are also included. After vigorous vortexing for 10 seconds, the mixture is left to react for 30 minutes in the dark at room temperature. The absorbance is measured at 517 nm using a spectrophotometer. The DPPH radical scavenging activity is calculated using the following formula: DPPH Free Radical Inhibition (%) = [1 - (Absorbance of the test sample) / (Absorbance of the control sample)] × 100

### 2.8. Recovery assessment against H_2_O_2_

During cell culturing, experiments were conducted using two sets: one for measuring cell viability and the other for staining with Methylene Blue solution. HaCaT cells were cultured in a 24-well plate, reaching approximately 90% confluence, and left to incubate for 24 hours. Subsequently, the cells were treated with samples and 1mM H2O2 for 12-16 hours. A positive control group with 2mM NAC was also included in the treatment. Three replicates were utilized to the CCK-8 assay for assessing cell viability, while the remaining three replicates were used for staining. After removing the medium from the 24-well plate, 500μL of 4% PFA solution per well was added for fixation at room temperature for 20 minutes. Following removal of the 4% PFA solution and PBS washing, the cells were stained using a 0.2% Methylene Blue solution. Post staining, images were captured using a microscope (Motic Asia, Hong Kong).

### 2.9. Minimum inhibitory concentration test (MIC assay)

For the MIC assay, Staphylococcus epidermidis, Staphylococcus aureus, and Escherichia coli strains were utilized. Bacterial stocks were thawed from -80°C and activated on a liquid medium. The bacterial suspension was adjusted to an OD600 of 0.1, and a maximum of 1% volume was inoculated into 5mL of liquid medium to prevent overgrowth. Two kinds of HCDP, WVYY, and PSLPA were treated respectively, with a positive control using 0.5 mg/mL Ampicillin. Following inoculation, bacterial cultures were incubated for 16 hours. Post-incubation, bacteria were serially diluted (10^1^ to 10^7^ dilutions) and mixed in dilution solutions of the microbial medium. Thorough vortexing ensured homogeneous mixing before aliquoting 100μL of appropriately diluted bacterial suspension onto solid agar plates for culturing. Colonies on completed solid agar plates were counted. Colony-forming units (CFUs) were determined by multiplying the colony count with the dilution factor. Colony counts were measured using the Scan 1200 (Interscience, France) equipment, and CFU calculations were performed using the formula: Bacterial count (%) = (Number of colonies in the test sample / Number of colonies in the control sample) x 100.

### 2.10. Melanin content assay

B16F1 cells, a mouse melanoma cell line, were plated at a density of 1×10^5^ cells per well in 6-well plates and cultured for 24 hours under standard cell culture conditions. Subsequently, the cells were treated with HSCE and continued to be cultured for an additional 72 hours. A positive control containing 100 ppm Kojic acid was included, and each sample received an additional 1 μM α-MSH to stimulate melanin production. Following the incubation period, the upper layer of the medium was removed, and cells were harvested using 1X Trypsin-EDTA. To dissolve melanin, a solution of 1 N NaOH (300 μL) was added to the cell pellet, and the mixture was boiled at 100°C for 30 minutes. The resulting dissolved melanin solution was partially transferred to a 96-well plate for absorbance measurement at 405 nm. The absorbance value for each sample was quantified relative to the protein content, and the melanin production inhibition rate was expressed as a percentage. The calculation for the melanin production inhibition rate is given by the formula: Melanin production inhibition rate (%) = [1 - (test group melanin amount / α-MSH treated control melanin amount)] * 100.

### 2.11. Measurement of intracellular glutathione (GSH) and reactive oxygen species (ROS) levels

Following HCDP treatment, the cells underwent thorough PBS washing. Next, intracellular ROS and GSH levels in keratinocytes were measured using H2DCFDA (2’, 7’- dichlorodihydrofluorescein diacetate; Invitrogen) and CellTracker Blue (4-chloromethyl- 6.8- difluoro- 7- hydroxycoumarin; CMF2HC; Invitrogen), respectively. For staining, cells were exposed to either 10 μM of CellTracker Blue or 10 μM of H2DCFDA in PBS and left to incubate for 30 minutes at room temperature while avoiding light. After staining, cells were washed again in PBS multiple times. Fluorescence intensities were measured using a fluorescence microscope (BX-53; Olympus, Japan) with UV filters (460 nm for ROS and 370 nm for GSH), followed by image capture. Image J software was utilized for analysis, standardizing the intensities of the control group to 1.

### 2.12. Detection of lipid peroxidation

Following the administration of HCDP, cells underwent a comprehensive PBS wash and were subsequently exposed to 5 μM of BODIPY™ 581/591 C11 (A12410; Molecular Probes, USA) in PBS for 1 hour at 37°C in a light-avoided environment. Upon staining, cells were once again washed with PBS and mounted on glass slides, covered with coverslips. For image acquisition, a fluorescence microscope (BX-53; Olympus, Japan) equipped with fluorescent filters (591nm for non-peroxidation and 510nm for peroxidation) was utilized. The ImageJ software (version 1.46r; National Institutes of Health, USA) was employed to measure lipid peroxidation intensities. The intensities of the control group were standardized to 1.

### 2.13. Immunofluorescence staining

Following treatment with HCDP, the cells were thoroughly washed with PBS. Subsequently, the cells were fixed using 4% paraformaldehyde (w/v) in PBS for 1 hour at room temperature. To permeabilize the cells, 1% Triton X-100 (v/v) in distilled water (DW) was applied for 1 hour at 37°C, followed by four washes in 1% PVA/PBS. To minimize nonspecific binding, the cells were then incubated in 2% BSA in 1% PVA/PBS for 2 hours. The cells were directly exposed to primary antibodies for NRF2 (1:200; PA5–34401; Thermo Fisher Scientific) and Heme Oxygenase 1 (HO-1) (1:200; ab13248; Abcam), and incubated overnight at 4°C. After this incubation, the cells underwent multiple washes in 1% PVA/PBS and were then treated with a secondary fluorescein isothiocyanate-conjugated anti-rabbit polyclonal antibody (1:200; ab6717; Abcam, Cambridge, UK) at 37°C for 2 hours while avoiding light. Following the secondary antibody incubation and subsequent washes with 1% PVA/PBS, the cells were immediately counterstained with 5 μg/mL Hoechst-33342 for 8 minutes. Post thorough washing, they were mounted on glass slides, covered with cover slips, and examined under a fluorescence microscope. The evaluation of fluorescence was conducted using ImageJ software (version 1.46r; National Institute of Health, USA). Evaluation of lipid peroxidation

### 2.14. Analysis of gene expression by quantitative real-time PCR

The cells treated with HSDP were detached using trypsin and washed with PBS before storage at -80°C until RNA extraction. For this purpose, a minimum of 1 x10^6^ cells per experimental group were processed using the RNeasy Mini kit (#74104; QIAGEN). The mRNA levels were measured with the NanoDrop (DS-11, DeNovix), followed by cDNA synthesis using RT Master Mix (FSQ-201, TOYOBO, Japan) following the manufacturer’s instructions. Quantitative real-time PCR (qRT-PCR) involved reaction mixtures containing 10 μL THUNDERBIRDTM Next SYBR® qPCR Mix (QPS-201, TOYOBO, Japan), 1 μL (10 pmol/μL) of forward primer, 1 μL (10 pmol/mL) of reverse primer, 7 μL of Nuclease-Free Water, and 1 μL of cDNA in a PCR plate. The amplification was carried out with the Rotor-Gene Q 6plex System (Rotor-Gene Q, Qiagen, Germany) through 40 cycles: denaturation at 95°C for 15 s, annealing at 62°C for 1 min, and extension at 72°C for 1 min. Each plate had at least three replications. The quantification of each target gene expression was standardized against the endogenous control gene (GAPDH) and a list of the primers is shown in Table S1. Relative expressions were calculated using the formula:

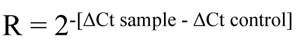

### 2.15. Statistical analysis

The SigmaStat statistical software (SPSS, Inc., Chicago, IL, USA) was utilized for thorough statistical analysis in this study. Each experiment was replicated a minimum of three times, covering both biological and technical replications. Results are depicted as means ± standard error of the mean. Before analysis, all data underwent normality and homoscedasticity assessments. To identify significant differences among three or more groups, the Kruskal– Wallis test (for non-normally distributed data) and one-way ANOVA (for normally distributed data) were performed. Post hoc analysis for one-way ANOVA entailed Duncan’s multiple range test (for equal variance) or Dunnett’s T3 test (for unequal variance). When comparing two groups, the Mann–Whitney U test (for non-normally distributed data) and Student’s t-test (for normally distributed data) were applied. A significance threshold of p < 0.05 was considered. Additionally, the data were reassessed, and visual representations were generated using GraphPad PRISM 5.01 (PRISM 5, GraphPad Software, USA).

## 3. Results

### 3.1. Optimization of the concentrations of the HSDPs PSLPA and WVYY concentrations

In the present study, we investigated the effects of two HSDPs WVYY and PSLPA. First, we determined their optimal concentrations in human keratinocytes (HaCaT) and fibroblasts (Detroit cells) using the CCK-8 assay. Both peptides were initially tested at 5, 10, 50, and 100 μM. Significant increase in the viability of fibroblasts was noted at 100 μM with WVYY treatment (p < 0.05), and at 50 μM (p < 0.05) and 100 μM (p < 0.001) with PSLPA treatment (Fig S1a). On the other hand, 10 μM of WVYY treatment (p < 0.05) and 50 μM of PSLPA treatment (p < 0.05) significantly increased the viability of keratinocytes (Sb1 Fig). To the best of our knowledge, this is the first report of the effect of these peptides on the viability of fibroblasts and keratinocytes. Based on the preliminary findings with the initial concentration ranges, we subsequently investigated the effect of 25, 50, 75, and 100 μM to determine the optimal peptide concentrations. Consistently, WVYY significantly increased the viability of fibroblasts at all tested concentrations (p < 0.0001), while PSLPA significantly increased the viability of fibroblasts at 25 and 75 μM (p < 0.05) (Fig 1a). Notably, an increase in the viability of PSLPA-treated keratinocytes was noted at 50 μM; however, the effect was not significant (Fig 1a). On the other hand, WVYY did not have an effect on the viability of keratinocytes at any tested concentration, whereas PSLPA significantly increased the viability of keratinocytes at all tested concentrations (p < 0.05; Fig 1b). Therefore, the optimized concentrations of WVYY and PSLPA were verified and confirmed Based on these findings, we selected 50 μM of PSLPA and 25 μM of WVYY as the optimal concentrations. Additionally, we explored the potential synergistic effects of 25 μM WVYY and 50 μM PSLPA in combination. Remarkably, all treatments (25 μM WVYY, 50 μM PSLPA, and the combination) significantly increased the viability of fibroblasts compared to that in the control group (p < 0.05). On the other hand, 25 μM WVYY and 50 μM PSLPA, and not the combination, significantly increased the viability of keratinocytes (p < 0.0001; Fig 1c). These results confirm the efficacy of both peptides as well as their combination at the determined optimal concentrations.

**Figure 1.**
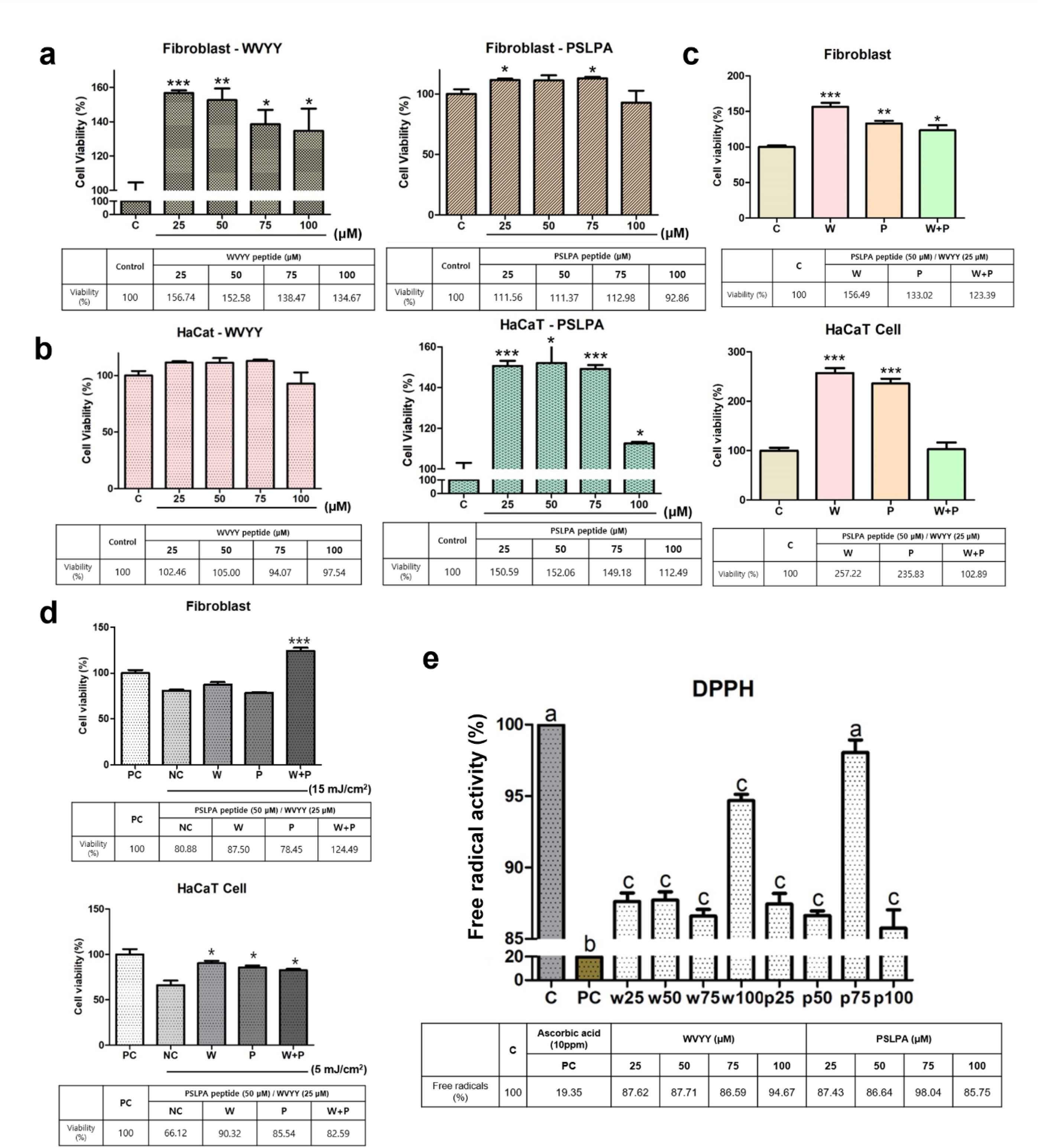
Optimization of WVYY and PSLPA in fibroblasts and keratinocytes. Each peptide was treated with fibroblasts and keratinocytes at concentrations of 25, 50, 75, and 100 μM to confirm cell viability. (a and b) Subsequently, the optimized concentrations were determined based on cell viability, including an experimental group with a mixture of the two peptides at their optimal concentrations, and the trend in cell viability at each optimal concentration was reconfirmed. (c) Additionally, the protective function against ultraviolet rays was evaluated when each peptide’s concentration and mixed treatment were applied. PC represents the untreated control with ultraviolet rays and reagents, while NC is the control treated only with ultraviolet rays without reagents. (d) Finally, to confirm if each peptide itself reacts to free radicals, the antioxidant effects were verified at all concentrations through DPPH. PC used 10 ppm Ascorbic acid. Significant differences in (a to d) were indicated with asterisks (p < 0.05 to 0.0001), and significant differences in (e) were indicated with alphabetical letters (p < 0.05). PC: Positive control; NC: Negative control; C: Control; W: WVYY; P: PSLPA.

### 3.2. HSDPs WVYY and PSLPA exhibit protective effects against ultraviolet radiation and free radicals

Next, we evaluated the protective abilities of the HSDPs WVYY and PSLPA against ultraviolet stimulation. To this end, we used a positive control (PC), i.e., cells that were not subjected to ultraviolet exposure, and a negative control (NC), i.e., cells that were subjected to ultraviolet exposure in the absence of peptide treatment. Interestingly, at 5 or 15 mJ/cm^2^, only treatment with the combination group exhibited significant protection against ultraviolet radiation exposure in fibroblasts, even surpassing the PC (p < 0.0001). On the other hand, all experimental groups significantly protected keratinocytes against ultraviolet radiation exposure, as evident by increased viability compared to that in the NC (p < 0.05; Fig 1d). These findings suggest that the two HSDPs possess ultraviolet protective properties, although further investigation is warranted. Given these findings, the free radical scavenging activity of the two HSDPs was assessed using DPPH, and ascorbic acid (10 ppm) was used as the PC. A significant decrease in free radical activity was observed at several concentrations with both HSDPs compared to that in the control (p < 0.05; Fig 1e). In summary, the two peptides themselves possess defensive properties against ultraviolet radiation and free radicals. Therefore, the final working concentrations of WVYY and PSLPA for subsequent analysis were determined through multiple approaches.

### 3.3. Asssessment of recovery and defense properties of WVYY and PSLPA

Given that we determined the optimal concentrations of the two peptides, we further examined various biological effects including effect on melanin content, antibacterial activity, effect on wound healing, and H_2_O_2_ defense capacity. No differences were observed in the melanin content in keratinocytes in any of the experimental groups, suggesting that the two peptides did not have any inhibitory effects on melanin synthesis (S1c Fig). For the assessment of antibacterial effects, we used three bacterial strains: *Escherichia coli*, *Staphylococcus aureus*, and *Staphylococcus epidermidis*. Minimum inhibitory concentration (MIC) assays were performed to assess the antibacterial effects and identify suitable concentrations for bacterial growth inhibition. Initially, liquid bacterial cultures were evaluated to measure the effect of the two HSDPs on colony-forming units by determining the OD 600. A significant reduction in the colony counts for *E. coli* was noted with 75 μM WVYY and 50 μM PSLPA (p < 0.0001 and p < 0.01, respectively; Fig 2a’). Similarly, a significant reduction was also noted in the colony counts for *S. aureus* with 75 μM WVYY and 75 μM PSLPA (both p < 0.001; Fig 2b’). However, reduction in *S. epidermidis* colony counts was not noted in any of the experimental groups (Fig 2c’). Thus, using the optimized concentrations, the antibacterial effect of the two HSDPs was reassessed. Consistently, we found that WVYY and PSLPA inhibited *E. coli* growth by 29.24% and 14.04% (Fig 2a and 2a’’), respectively. Similarly, WVYY and PSLPA inhibited *S. aureus* growth by 48.81% and 35.63%, respectively (Fig 2b and 2b’’). Interestingly, and in contrast to *E. coli* and *S. aureus,* stimulation of *S. epidermidis* growth was noted with both WVYY and PSLPA, with inhibition rates of −47.70% and −81.18%, respectively (Fig 2c and 2c’’). Therefore, it has been demonstrated that WVYY and PSLPA exhibit inhibitory functions against harmful bacteria and even contribute to the proliferation of beneficial bacteria, such as Staphylococcus epidermidis.

**Figure 2.**
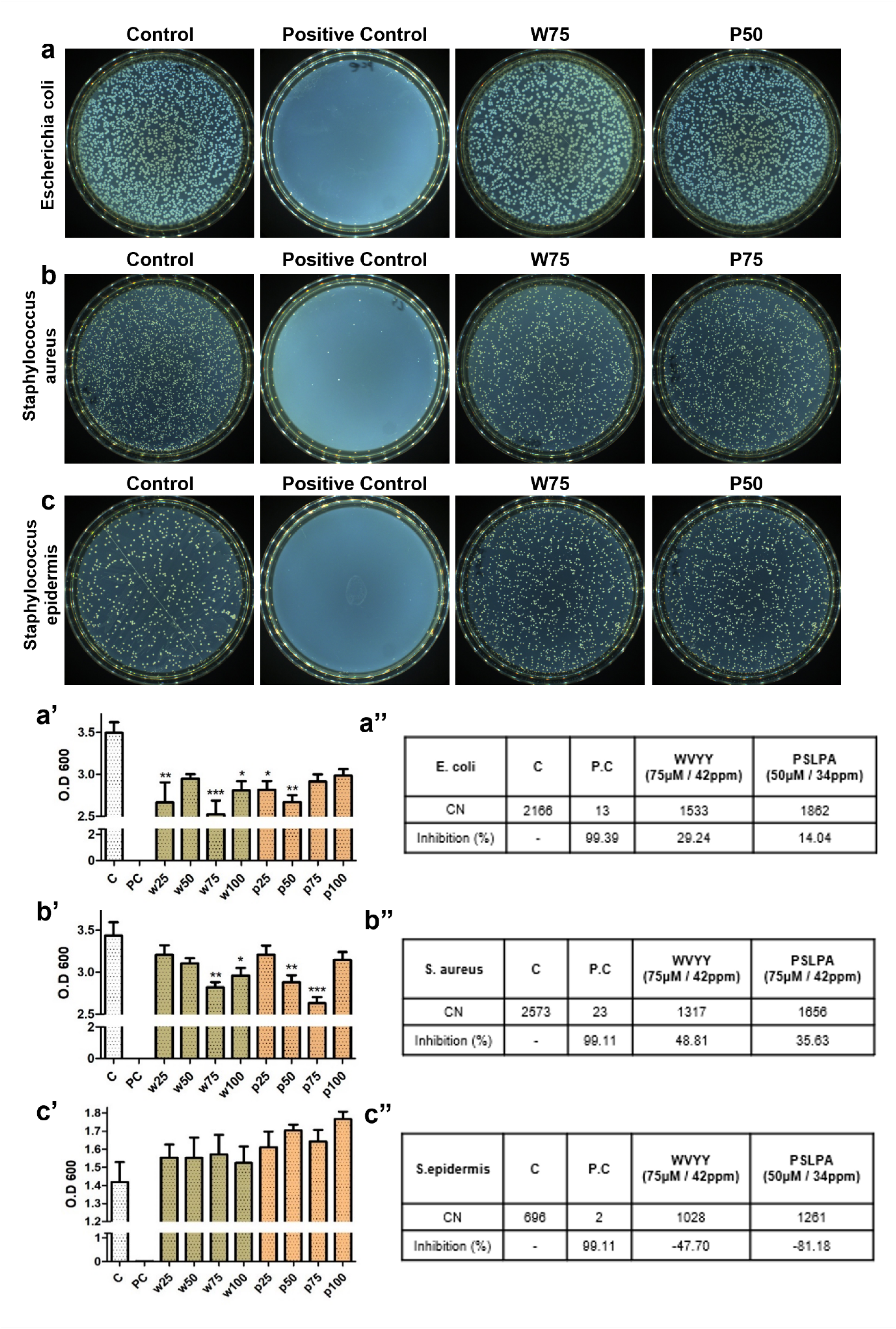
The antibacterial test was conducted on Escherichia coli, Streptococcus aureus, and Streptococcus epidermidis (a-c). Serial dilutions were performed through MIC testing, measuring bacterial growth in terms of O.D values. The obtained values were then plated on solid media, and colony counts were measured (a’-c’’). By selecting the concentration at which the number of bacterial colonies significantly decreased, the inhibition rate was determined. PC used 0.5 mg/ml Ampicillin. All significant differences were denoted with asterisks (p < 0.05 to 0.0001). PC: Positive control; CN: Colony number; C: Control; W: WVYY; P: PSLPA.

Next, the effect of the two HSDPs on wound healing was evaluated using keratinocytes. Significant wound healing was noted across all groups, WVYY, PSLPA, and their combination (p < 0.05; Fig 3b). Although a statistical comparison was not performed, when calculating the area at the pixel level, WVYY showed 38.10% compared to T0, PSLPA showed 43.63%, and the combination treatment showed 66.70%. Furthermore, when compared to T18, WVYY showed 20.52%, PSLPA showed 26.05%, and the combination treatment showed 49.12%. Lastly, antioxidant effects of the two HSDPs against H2O2 were assessed. A significant defense against free radicals in cell viability for the PSLPA alone (85.3%) and the combination of WVYY and PSLPA (85.85%) significantly increased the viability of H2O2-treated cells (p < 0.05; Fig 3c, 3c’, and 3d), suggesting antioxidant effects; however, the same was not observed with WVYY (73.72%). Collectively, these findings suggest that both HSDPs demonstrate antibacterial, wound-healing, and antioxidative effects.

**Figure 3.**
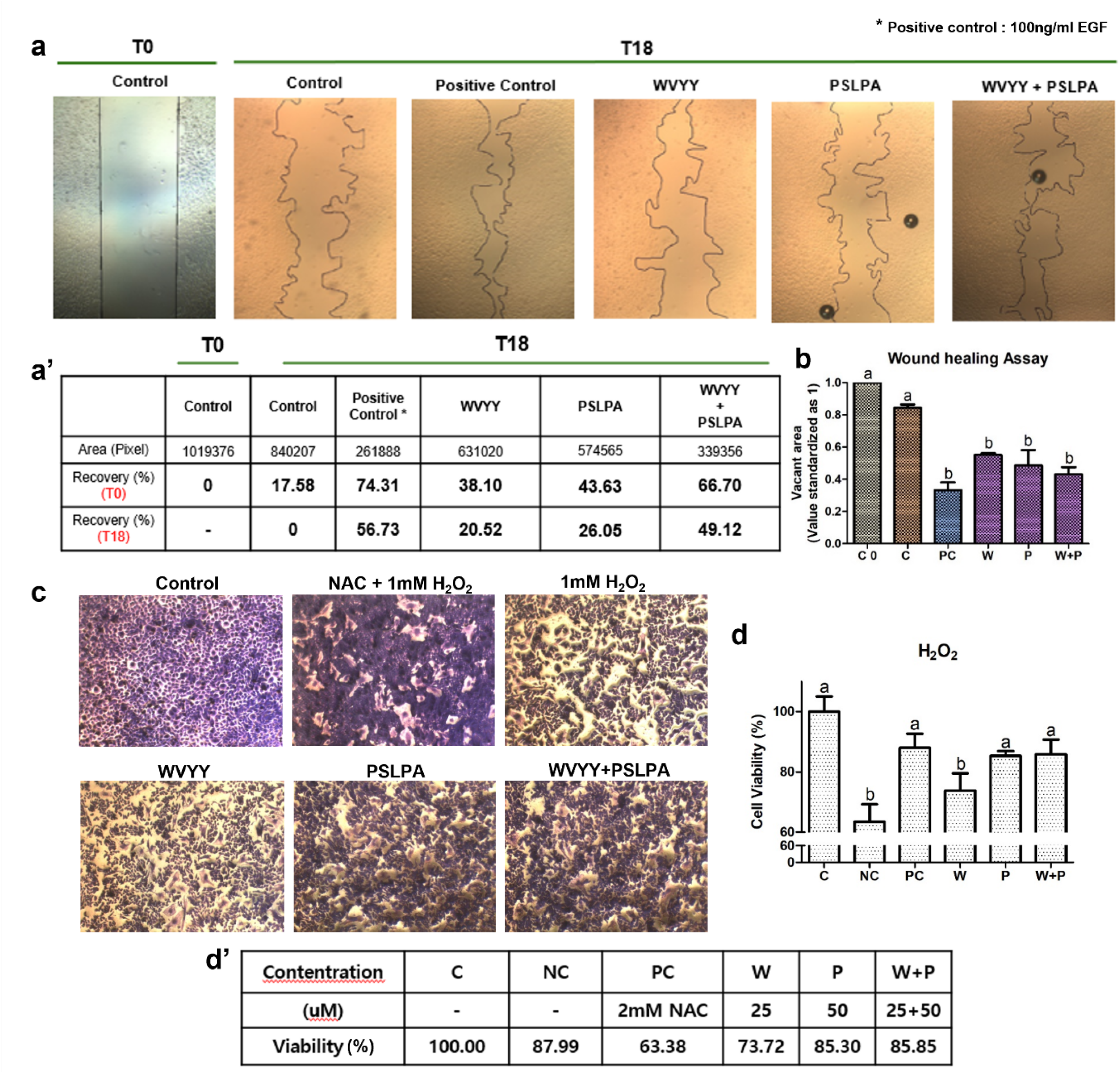
The wound healing and antioxidant tests of WVYY and PSLPA were conducted on keratinocytes using the optimal concentrations and mixed treatment groups determined in the previous experiment. (a) Keratinocytes were seeded, and wound healing was initiated at T0. After 18 hours (T18), the healed area was measured to evaluate the extent of recovery. PC was treated with 100 mg/ml EGF for comparison. (a’ and b) Simultaneously, the antioxidant effects of WVYY and PSLPA were assessed by treating with H2O2 to observe the effects of each peptide individually and in combination on free radicals. After staining with methylene blue, the antioxidant effects were visually confirmed (c), and cell viability was reconfirmed. PC was treated with NAC 1mM and H2O2 together. (d and d’) All significant differences were indicated with alphabetical letters (p < 0.05). T0: 0 Hours of wound; T18: 18 hours of wound; PC: Positive control; NC: Negative control; EGF: Epidermal growth factor; NAC: N-Acetyl Cysteine; C: Control; W: WVYY; P: PSLPA.

### 3.4. The HSDPs WVYY and PSLPA modulate the Nrf2 signaling pathway

Morphological validation confirmed the efficacy of both peptides and next, we investigated the mechanism underlying the observed biological effects of the two HSDPs. To this end, the Nrf2 signaling pathway, which is associated with antioxidant mechanisms, was examined using immunocytochemistry (ICC). Consistently, the expression of HO-1, one of the downstream targets of Nrf2, was evaluated. A gradual and significant increase in HO-1 expression was noted in all experimental groups (p < 0.05; Fig 4a and 4b). Moreover, the activation of Nrf2 was found to be significantly higher in the PSLPA and combination groups compared to that in the control group (p < 0.05; Fig 4c and 4d). Collectively, these findings suggest that both HSDPs, either individually or in combination, modulated the Nrf2 signaling pathway.

**Figure 4.**
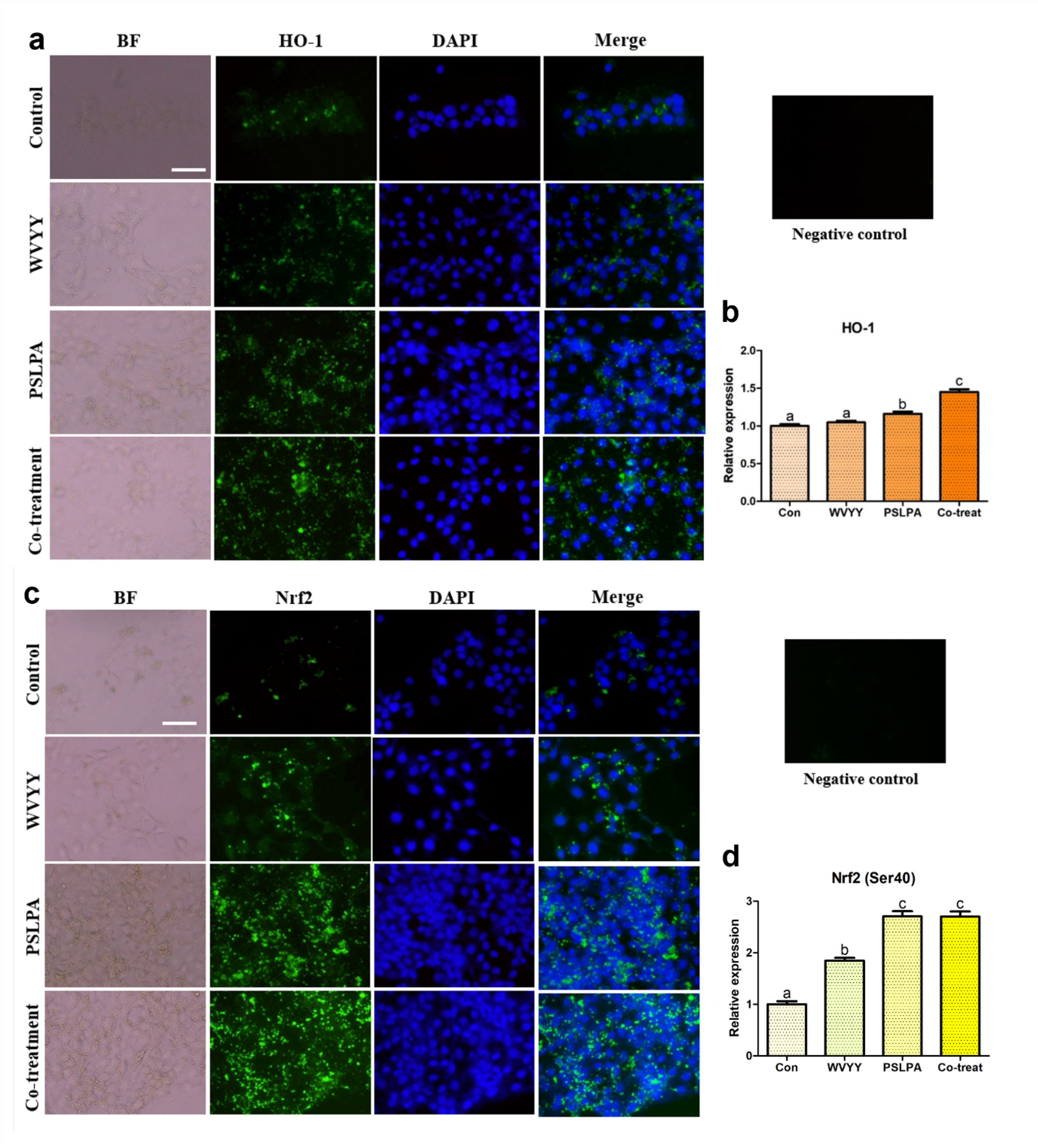
Immunocytochemistry was employed to assess the expression of HO-1 and Nrf2 proteins in keratinocytes. Nrf2, a pivotal factor in the Nrf2 signaling pathway, was examined in representative forms (a and b), while its downstream antioxidant responsive element, HO-1, was evaluated (c and d). The assessments were conducted in response to treatments with optimized concentrations of WVYY, PSLPA, and the combined treatment of both peptides. The intensities of the target proteins were measured, and normalization was performed using the intensity of the negative control, which was assessed in the absence of the primary antibody. This normalization strategy ensured uniform exposure times across all experimental groups. All significant differences were indicated with alphabetical letters (p < 0.05). White bar: 50 μm. Con: Control; W: WVYY; P: PSLPA; Co-treatment: WVYY+PSLPA; BF: Bright Field

The following investigated the closely related processes of Nrf2 signaling, lipid peroxidation, and the levels of ROS and GSH. Significantly higher levels of non-peroxidized lipids and consistently, significantly lower levels of peroxidized lipids were observed with both PSLPA and WVYY treatment compared with that in the control group (all p < 0.05; Fig 5). Remarkably, similar findings were also noted with the combination group with significantly higher levels of non-peroxidized lipids and significantly lower levels of peroxidized lipids compared with that in the control group (p < 0.05; Fig 5a–5c). When comparing the ratio of non-peroxidized and peroxidized lipids, the PSLPA and combination groups exhibited the most significant differences (P < 0.05; Fig 5d). Next, GSH and ROS levels were evaluated. Significantly lower ROS levels were noted in all experimental groups compared to that in the control group (all p < 0.05; Fig 5e). On the other hand, only the PSLPA and combination groups exhibited a significant increase in GSH expression compared to that in the control group (p < 0.05; Fig 5e). These findings substantiate the antioxidant effects of the two HSDPs at the protein level, consistent with the ICC findings.

**Figure 5.**
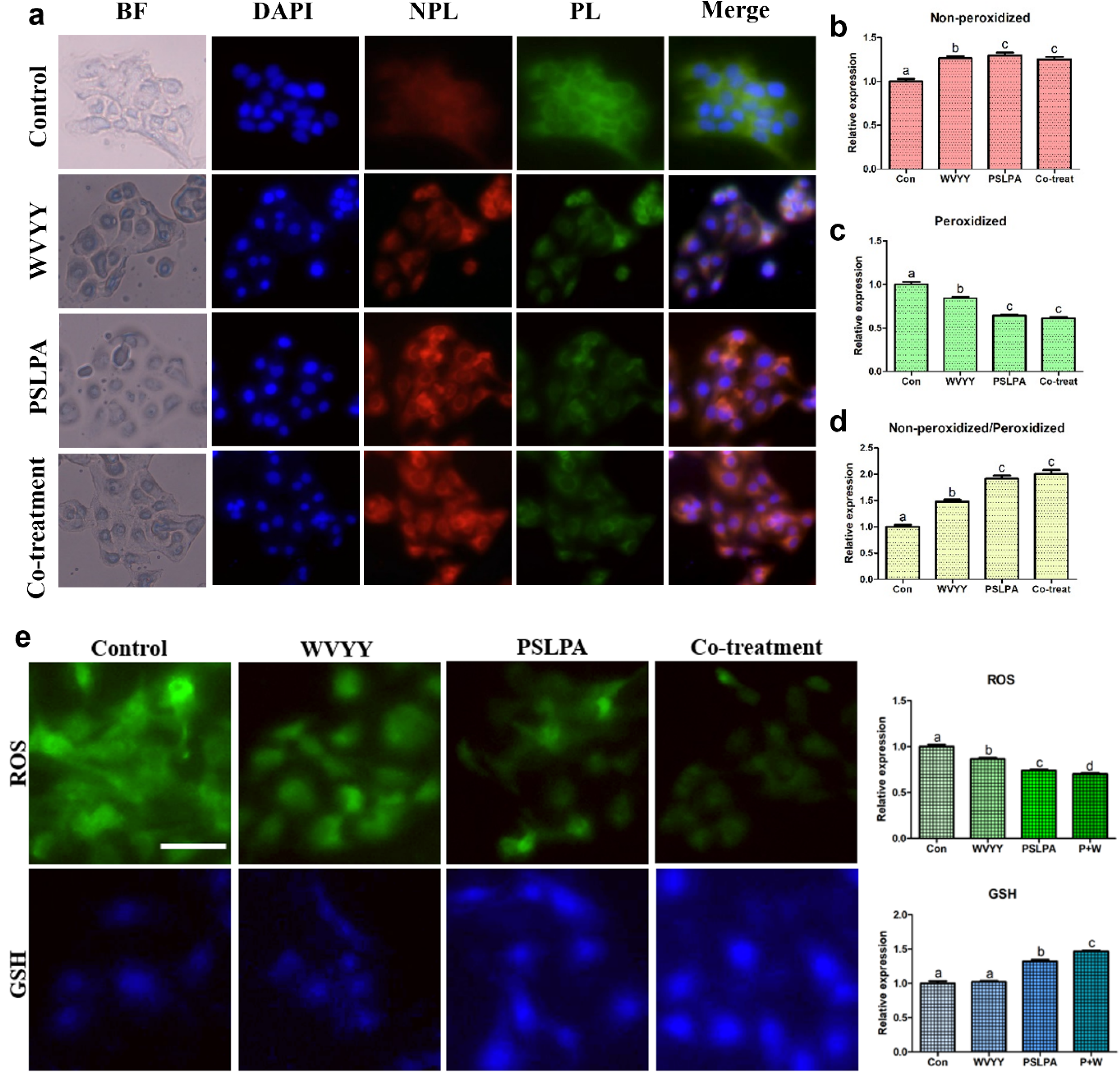
BODIPY staining for lipid peroxidation and ROS/GSH staining were conducted in keratinocytes following treatments with WVYY, PSLPA, and the co-treatment of both peptides. For non-peroxidized samples, excitation and emission wavelengths were in red, measuring 581/591 nm, while peroxidized samples exhibited green fluorescence, with excitation and emission at 488/510 nm. (a-c) The merged images of DAPI, non-peroxidized, and peroxidized samples were analyzed to confirm the extent of expression differences, presented as ratios for each. (d) Furthermore, keratinocytes were cultured, and ROS and GSH were stained separately to measure oxidative stress levels and GSH production rates (e). All significant differences were denoted with alphabetical letters (p < 0.05). The scale bar represents 50 μm. Con: Control; W: WVYY; P: PSLPA; Co-treatment: WVYY+PSLPA; BF: Bright field; ROS: Reactive oxygen species (ROS), GSH: Glutathione; NPL: Non-peroxidized; PL: Peroxidized.

Further, qRT-PCR was performed to confirm the involvement of the Nrf2 signaling pathway, and to assess the expression of antioxidant genes, keratinocyte growth factor (KGF), and apoptosis-related genes. With respect to the Nrf2 signaling pathway-related genes, all experimental groups exhibited a significant increase in *Nrf2* expression compared to that in the control, with the combination group showing the highest expression (p < 0.05; Fig 6a). On the other hand, significantly lower expression of *Keap1* expression was noted in the PSLPA group compared to that in the control group (p < 0.05; Fig 6a). Whereas, *Cul3* expression was significantly higher in the combination group compared to that in the control group (p < 0.05; Fig 6a). Among the antioxidative mechanism-related genes a significant increase in *HO-1* expression was noted in all experimental groups compared to that in the control group, with the combination group exhibiting the highest expression (p < 0.05; Fig 6b). *CAT* expression was also significantly increased in all experimental groups compared to that in the control group (p < 0.05; Fig 6b), with the WVYY group exhibiting the highest expression. *SOD1* expression was significantly higher in the PSLPA group compared to that in the control group (p < 0.05), whereas *SOD2* expression was significantly higher in the PSLPA and combination groups compared to that in the control group (p < 0.05; Fig 6b). *KGF* expression was significantly higher in the PSLPA group compared to that in the control group (P < 0.05; Fig 6c), and similar trends were observed in the other experimental groups.

**Figure 6.**
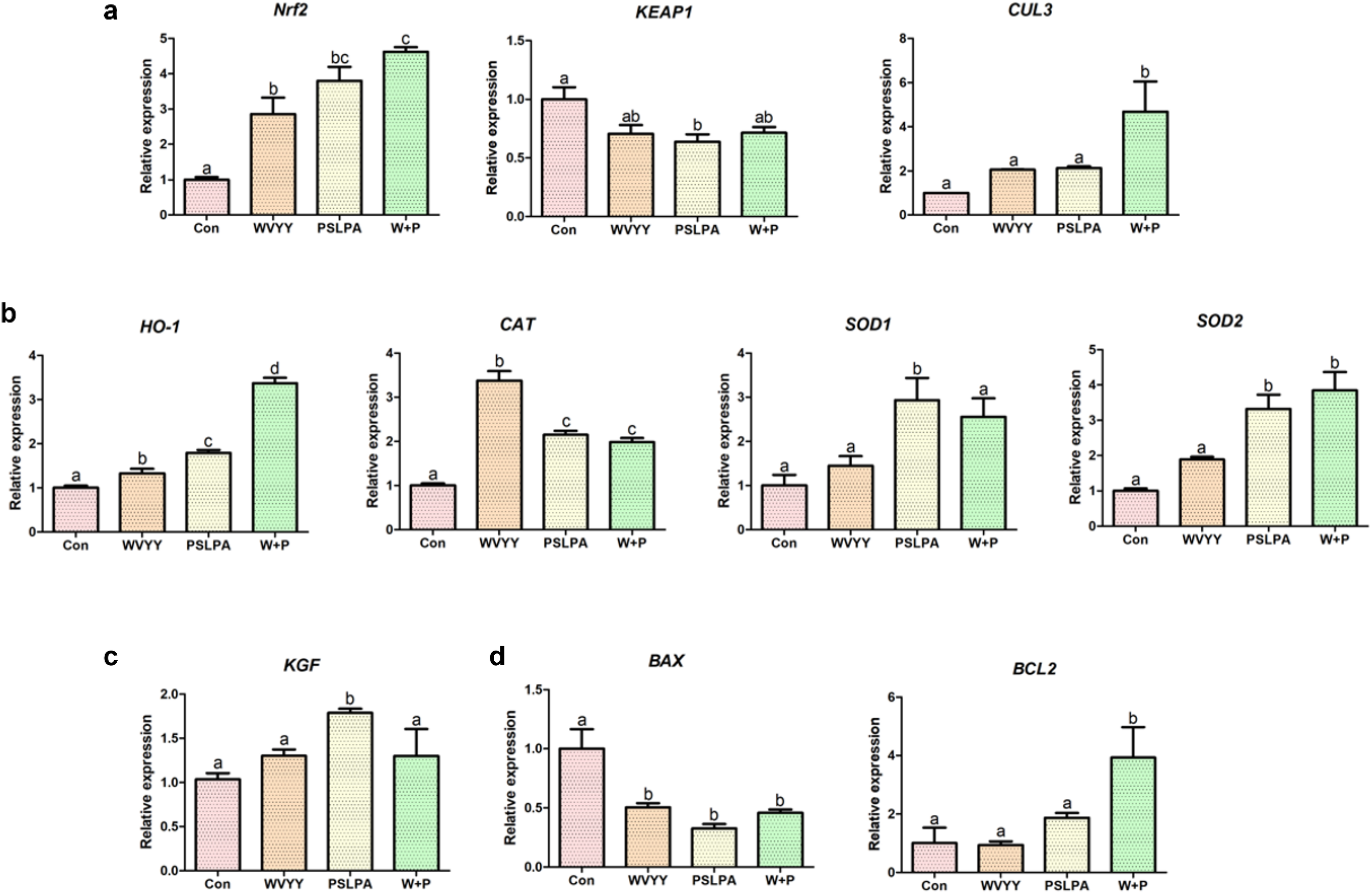
Analysis of the relative expression of genes associated with (a) the Nrf2 signaling pathway (*Nrf2*, *KEAP1*, and *CUL3*), (b) antioxidant responsive elements (*HO-1*, *CAT*, *SOD1*, and *SOD2*), (c) cell proliferation (*KGF*), and (d) apoptosis (*BAX* and *BCL2*) was performed using quantitative reverse transcriptase-polymerase chain reaction (qRT-PCR). Triplicates of technical and biological replications were conducted. Significant differences were denoted by alphabetical letters (p < 0.05). Con: Control; W: WVYY; P: PSLPA; Co-treatment: WVYY+PSLPA.

Finally, with respect to apoptosis-related genes, *B-cell lymphoma 2* (*BCL2*) expression was significantly decreased only in the combination group compared to that in the control group (p < 0.05; Fig 6d), whereas *Bcl-2-associated X protein* (*BAX*) expression was significantly lower expression in all experimental groups compared to that in the control group (p < 0.05; Fig 6d). Collectively, these findings provide further evidence to the antioxidant effects of the HSDPs WVYY and PSLPA, and confirm the involvement of the Nrf2 signaling pathway in the same.

## Discussion

In the present study, we investigated the biological effects of two HSDPs. Our study addresses an important gap in the existing knowledge of the mechanism underlying the reported antioxidant effects of hemp products, including HSDPs. We considered that the effects of various hemp extracts could be complex, and among the many bioactive components in the extracts, we hypothesized that certain peptides might play a direct role in the biological effects, specifically in the antioxidant, wound healing, and antibacterial effects. Consistently, we confirmed that the HSDPs WVYY and PSLPA exhibited antioxidant and antibacterial effects and promoted wound healing. Jeshionek et al. reported that bioactive compounds, such as quercetin, luteolin, caffeic acid, and other secondary metabolites, can be detected in plant extracts [37]. Additionally, plant extracts can also contain other molecules, such as protein hydrolysate or bioactive peptides [38, 39]. Hence, while various substances derived from plant extracts can exert diverse biological effects, it is crucial to precisely identify the specific substances that exert the observed effects to exploit the same for other applications. In this regard, our findings demonstrate that HSDPs are involved in the antioxidant effects of hemp seed extracts.

We investigated the effects of HSDPs on wound healing and viability using keratinocytes. We also investigated their antioxidant effects using assays, such as H_2_O_2_ defense and DPPH. Additionally, we assessed their antibacterial effects against various bacterial species. The Nrf2 signaling pathway was specifically investigated in this study given its roles in antioxidant mechanisms, wound healing, and immune response to bacterial infections {Ahmed, 2017 #266;Kaspar, 2009 #263;Kim, 2020 #254;Li, 2021 #269}. We observed that the two HSDPs exerted antioxidant and antibacterial effects and promoted wound healing. Especially concerning bacteria, harmful bacteria such as E.coli and S. aureus decreased, while beneficial bacteria like S. epidermidis significantly increased. These results suggest that the roles of these two peptides may be somewhat crucial to skin microbiome, even from a cosmetic perspective. Given the role of the Nrf2 signaling pathway in the same, we investigated whether the two HSDPs modulate this pathway to exert their effects.

As mentioned previously, the Nrf2 signaling pathway plays a central role in mitigating oxidative stress through its regulation of antioxidant mechanisms [40]. During oxidative stress, Nrf2 is preferentially translocated into the nucleus following inactivation of Keap1. Subsequently, it dimerizes with sMaf to facilitate the transcription of antioxidant responsive element-containing genes, such as *SOD1*, *SOD2*, *HO-1*, and *CAT,* to mediate the antioxidant mechanism [41, 42]. Additionally, Nrf2 regulates the expression of enzymes involved in GSH production, quinone detoxification, and NADPH synthesis [43]. Moreover, the Nrf2 signaling pathway is involved in inflammation [34, 44], cell viability [45], antibacterial activity[36, 46], and wound healing [47, 48]. Specifically, Lwin et al. reported that the expression of epidermal growth factor, fibroblast growth factor, and vascular endothelial growth factor as well as that of Nrf2 are enhanced during wound healing of the keratinocytes. This alleviated inflammation and oxidative stress at the wound site, leading to the promotion of collagen deposition and angiogenesis [49]. Therefore, the Nrf2 signaling pathway is an important pathway in the antioxidant response.

Given the above-mentioned significance of the Nrf2 signaling pathway, we investigated its role in the biological effects mediated by WVYY and PSLPA. The involvement of the Nrf2 signaling pathway in the antioxidant effects of WVYY and PSLPA was confirmed by ICC and qRT-PCR. Specifically, when examining the expression of antioxidant-related genes (Fig 6b), distinct gene expression patterns were observed with WVYY and PSLPA treatment. For instance, *HO-1* expression was significantly higher in the WVYY and PSLPA groups compared to that in the control group, and synergistic effects were observed with combination treatment (Fig 4a and Fig 6b). In contrast, *CAT* expression was significantly higher expression in all experimental groups, with the highest expression in the WVYY group. Similarly, the expression of *SOD1* and *SOD2* expression was increased in the PSLPA and combination groups. Although both WVYY and PSLPA have been reported to exert antioxidant effects, the underlying mechanisms seem to be distinct. Nonetheless, the significant increase in *Nrf2* expression observed in all experimental groups (Fig 6a) confirms both WVYY and PSLPA induce their antioxidant effects through the Nrf2 signaling pathway. However, the mechanisms downstream of Nrf2 activation appear to be different as suggest by our data.

Additionally, increased GSH levels and decreased ROS levels, correlating with the involvement of the Nrf2 pathway were noted. Furthermore, *KGF* expression was significantly increased in PSLPA-treated cells, suggesting its role in PSLPA induced effects. Therefore, although both HSDPs contribute to the antioxidant effects of hemp seed extracts, the mechanisms underlying these effects maybe distinct, as seen in the present study.

Ferroptosis is a type of Fe^2+^-dependent cell death that was discovered relatively recently [50]. This mechanism is primarily induced by the accumulation of lipid peroxides. ROS are continuously generated through a mechanism known as the free radical chain reaction, which results in lipid peroxidation, damaging the cell membrane [51]. Specifically, polyunsaturated fatty acids such as linoleic and arachidonic acids are involved in this process [52]. Dodson et al. reported that the Nrf2 pathways plays a crucial role in lipid peroxidation and mitigates the same [53]. Consistently, other studies have reported that the Nrf2 signaling pathway inhibits lipid peroxidation and ferroptosis [54–56]. Additionally, some reports have suggested a relationship between ferroptosis and GSH [57, 58]. Our findings further support the claims of the aforementioned studies, as we observed that the expression of GSH and Nrf2 and lipid peroxidation levels were modulated by the two HSDPs. Further, the connection of both peptides to each mechanism is evident, and as shown in Figure 5a–5d, PSLPA had a significantly greater effect on lipid peroxidation than WVYY.

## Conclusion

Previous studies have indicated the involvement of the Nrf2 signaling pathway in antioxidant mechanisms, wound healing, immune response to bacterial infections, and lipid peroxidation. In the present study, we confirmed the role of the Nrf2 signaling pathway in the observed effects of the HSDPs WVYY and PSLPA, including antioxidant and antibacterial effects, and wound healing. Further, distinct mechanisms were involved in the antioxidant effects of the two HSDPs. The key aspect of our study lies in understanding the varied roles of HSDPs in antioxidant mechanisms, and more detailed research is needed to precisely elucidate the mechanisms underlying the effects of both peptides.

Although a more in-depth investigation into the precise roles of each peptide is necessary, we believe that a thorough examination of the highly specific roles of various peptides could lead to a broad range of medical and biological applications. In conclusion, we confirmed that the peptides WVYY and PSLPA derived from hemp seed extracts exhibit multiple effects, including antioxidant mechanisms. We propose that the overall "effect" of hemp extract originates from these contributions.

## Acknowledgements and funding

This work was supported by Advanced Technology Center Plus (ATC+, 20017936, Development of growth factors and antibody drugs using plant cell-based platform technology and global sales expansion of fragrance materials.) funded By the Ministry of Trade, Industry and Energy (MOTIE, Korea). We would like to thank Editage (www.editage.co.kr) for English language editing and Ye Eun Kim for figure arrangement.

## Author Contributions

Conceptualization: Hyo Hyun Seo

Data curation: Euihyun Kim

Formal analysis: Euihyun Kim

Funding acquisition: Jung Hun Lee, Sang Hyun Moh h

Investigation: Hyo Hyun Seo, Euihyun Kim

Methodology: Euihyun Kim, Jihyeon Jang

Project administration: Jung Hun Lee, Sang Hyun Moh

Resources: Hyo Hyun Seo

Software: Euihyun Kim Supervision: Sang Hyun Moh

Validation: Euihyun Kim, Jihyeon Jang Visualization: Euihyun Kim

Writing (Original draft)– Euihyun Kim Writing (review and Editing)– All authors

## Data Availability Statement

All relevant data are within the paper.

---

**Figure s1.**
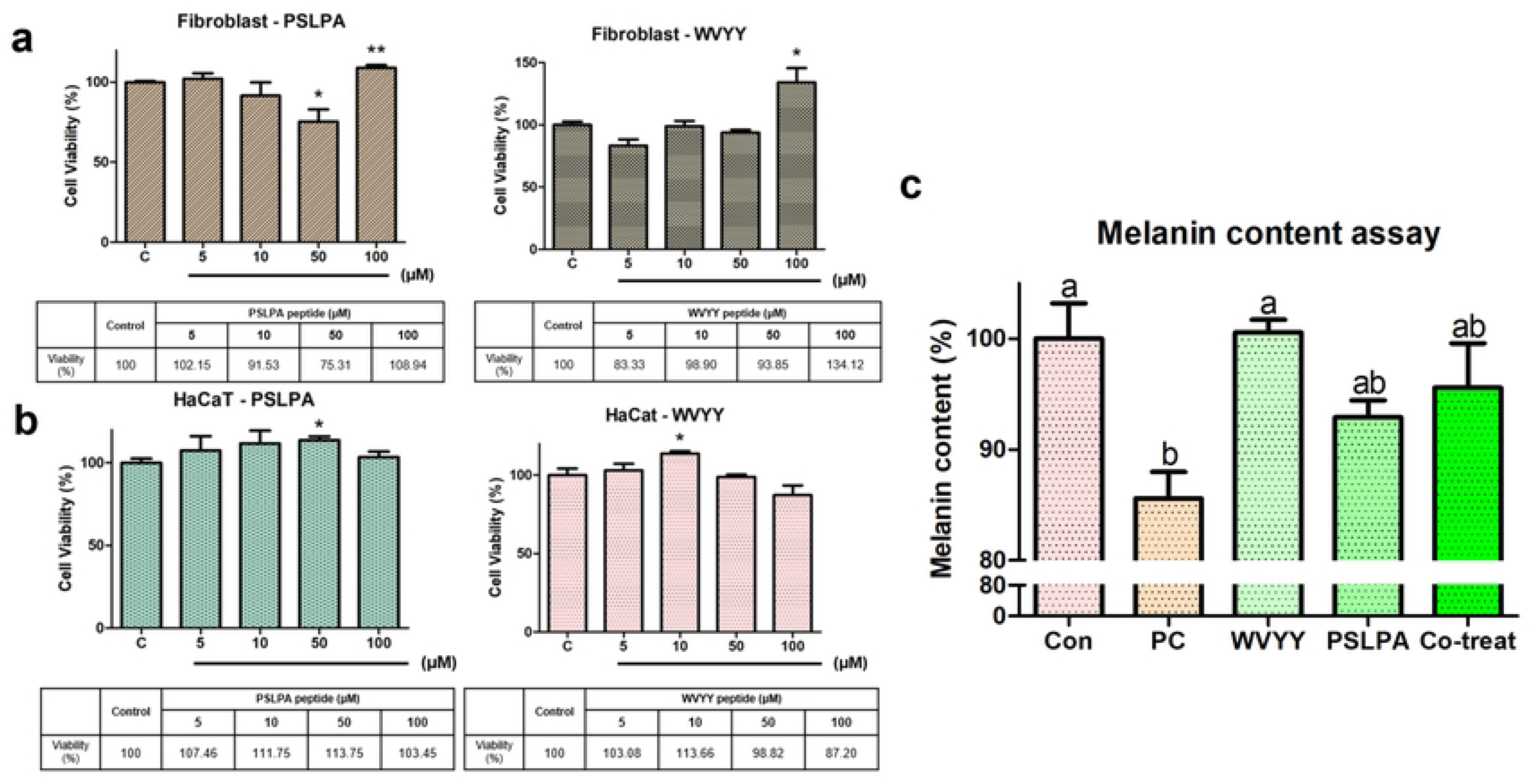

